# HSP90 inhibitors reduce cholesterol storage in Niemann-Pick type C1 mutant fibroblasts

**DOI:** 10.1101/2021.04.22.440982

**Authors:** Nina H. Pipalia, Syed Z. Saad, Kanagaraj Subramanian, Abigail Cross, Aisha al-Motawa, Kunal Garg, Brian S.J. Blagg, Len Neckers, Paul Helquist, Olaf Wiest, Daniel S. Ory, Frederick R. Maxfield

## Abstract

Niemann Pick type C1 (NPC1) disease is a lysosomal lipid storage disorder caused by mutations of the *NPC1* gene. More than 300 disease-associated mutations are reported in patients, resulting in abnormal accumulation of unesterified cholesterol, glycosphingolipids and other lipids in late endosomes and lysosomes (LE/Ly) of many cell types. Previously, we showed that treatment of many different *NPC1* mutant fibroblasts with histone deacetylase inhibitors resulted in reduction of cholesterol storage, and we found that this was associated with enhanced exit of the NPC1 protein from the endoplasmic reticulum and delivery to LE/Ly. This suggested that histone deacetylase inhibitors may work through changes in protein chaperones to enhance the folding of NPC1 mutants, allowing them to be delivered to LE/Ly. In this study we evaluated the effect of several HSP90 inhibitors on NPC1^I1061T^ skin fibroblasts. We found that HSP90 inhibition resulted in clearance of cholesterol from LE/Ly, and this was associated with enhanced delivery of the mutant NPC1^I1061T^ protein to LE/Ly. We also observed that inhibition of HSP90 increased the expression of HSP70, and overexpression of HSP70 also reduced cholesterol storage in *NPC1^I1061T^* fibroblasts. However, we did not see correction of cholesterol storage by arimoclomol, a drug that is reported to increase HSP70 expression, at doses up to 0.5 mM. These results indicate that manipulation of molecular chaperones may lead to effective treatments for NPC1 disease, but further investigation of mechanisms will be required.

## Introduction

Niemann-Pick type C1 (NPC1) disease is a neurovisceral lysosomal storage disease caused by mutations in both alleles of the *NPC1* gene. The incidence of occurrence in juveniles is estimated to be about one in 92,000, but with inclusion of mutations associated with late-onset disease the frequency increases to one in 20,000-36,000 (1). Over 300 disease-associated mutations reported (1–6), and biallelic mutations lead to abnormal accumulation of unesterified cholesterol, glycosphingolipids and other lipids in late endosomes and lysosomes (LE/Ly) of many cell types. The most common mutation, I1061T, is present in at least one allele of about 20% of patients (7). The clinical spectrum of NPC1 disease is very broad, ranging from a rapidly progressing fatal neonatal disorder to adult-onset neurodegeneration (8–10). NPC1 is a lysosomal glycoprotein with 13-transmembrane domains that is synthesized and folded with the aid of chaperones in the ER (11–14), where it acquires 14 N-linked glycans (15, 16). After exit from the ER, the NPC1 protein is exported to the Golgi and subsequently targeted to LE/Ly compartments (17). Many mutations in NPC1 result in protein misfolding and impaired trafficking of the protein out of ER, followed by protein degradation (ERAD) (11, 18). This results in significant reduction of NPC1 protein levels compared to wild type (WT) cells (11, 18). Lack of sufficient NPC1 protein in the LE/Ly leads to impaired cholesterol exit from these organelles.

There is no FDA-approved therapy to treat NPC1 disease. Intrathecal delivery of 2-hydroxypropyl beta-cyclodextrin (HPCD) was shown to slow disease progression in a phase 1/2a trial (19, 20), prompting a phase 2b/3 trial. A limitation of HPCD is that it does not cross the blood-brain barrier, requiring direct injection into the CNS. In a search for an orally available treatment, we showed that histone deacetylase inhibitor (HDACi) treatment of skin fibroblasts results in increased expression of NPC1 protein in mutant cells due to enhanced folding and exit from the ER (21). Using an engineered human cell line, we also observed that broad spectrum HDAC inhibitors can correct the NPC1 cholesterol storage defect in 60 of the 81 NPC1 mutants tested (18). These studies indicated that many of the mutant proteins could function adequately if they were delivered to LE/Ly. However, a study in a mouse model of *NPC1^I1061T^* (22) indicated that an HDACi (Vorinostat) provided no benefit in a whole animal model (23). Nevertheless, the correction in cell culture indicated the HDACi were assisting in correction of the disease phenotype for many *NPC1* mutations. In order to understand the molecular mechanism of HDACi on NPC1, we considered potential cellular effects of HDACi treatment. It has been shown that the NPC1^I1061T^ mutant protein undergoes ERAD, presumably associated with the inability to fold efficiently in the ER (11). Boosting chaperone activity has been proposed as a method to correct the defect in NPC1 mutant cells (12, 24–26). Changes in molecular chaperone activity are one of the many consequences of HDACi treatment (14, 27, 28). A quantitative proteomic analysis of the response to HDACi in NPC1 human fibroblasts revealed that the expression of proteins associated with protein folding was modulated (29). A detailed study of the response of many NPC1 mutations to allosteric modulators showed that many of the mutations were rescued to some degree by these modulators, which alter the interactions of HSP70 with co-chaperones (14). It has been shown that HSP90 as well as HSP70 can play a role in stabilization and correct folding of NPC1 (12).

Several HSP90 inhibitors have gone through extensive clinical testing as potential cancer therapeutics (30). We tested the effect of HSP90 inhibitors on the expression, folding, and activity of NPC1 in human fibroblasts that were homozygous for the *NPC1^I1061T^* allele, expecting that this would exacerbate the phenotype. Surprisingly, we found that HSP90 inhibition aided in transporting mutant NPC1 protein from the ER to LE/Ly, which led to increased clearance of cholesterol from LE/Ly. Molecular chaperones form a complex network to assist in protein folding in cells (31). HSP70 family members are key members of the chaperone family, and we confirmed that HSP90 inhibition led to an increase in HSP70 expression as reported previously (32, 33). Additionally, transfection of cells with HSP70-1A (HSPA1A) led to a reduction in cholesterol storage in *NPC1^I1061T^* cells. We tested arimoclomol, which is reported to increase expression of HSP70 and other chaperones (34), but we did not see an increase in HSP70 expression or a reduction in cholesterol storage in *NPC1^I1061T^* cells.

## Materials and methods

### Reagents

Gibco^®^ McCoy’s 5A Medium, Modified Eagle’s Medium (MEM), fetal bovine serum (FBS), Hank’s Balanced Salt Solution (HBSS), penicillin/streptomycin (P/S), Geneticin (G418), AlexaFluor-546 *N*-Hydroxy Succinimidyl Ester, and AlexaFluor 546 Goat anti-rabbit and AlexaFluor 488 Goat anti-Rat antibody, were purchased from Invitrogen Life Technologies Corporation (Carlsbad, CA). HSP90 inhibitors were from various suppliers: 17-AAG (Geldanamycin) from Reagent Direct (Encinitas, CA), AUY922 (Luminespib), Ganetespib and AT13387 (Onalespib) from Selleckchem, (Houston, TX) or from Blagg Laboratory (35), SNX2112 from Esanex (Indianapolis, IN) and TAS-116 from Taiho Pharmaceuticals (Tokyo, Japan), arimoclomol maleate (arimoclomol) was from Medkoo Biosciences, USA. GRP inhibitors were obtained from the Blagg laboratory and prepared as described in references in Table 1. All compounds were dissolved at 10 or 1 mM in dimethyl sulfoxide (DMSO) and stored at - 20°C. Acetylated low-density lipoprotein (AcLDL) was prepared by acetylation of LDL with acetic anhydride (36). AlexaFluor-546 labeled human LDL (LDL-A546) was prepared as described (37, 38). Rabbit polyclonal anti-human NPC1 (p-Rab anti-hNPC1) antibody was purchased from Abcam (Cambridge, MA), rabbit polyclonal anti-HSP40 (p-Rab anti-HSP40), anti-HSP70 (p-Rab anti-HSP70) and anti-HSP90 (p-Rab anti-HSP90) were purchased from Cell Signaling (Danvers, MA). A p-Rab anti-hNPC1 antibody was used for endo-H Western blotting assay. All other chemicals, including (DMSO, 99% fatty-acid free bovine serum albumin (BSA), filipin, paraformaldehyde (PFA) and 4-(2-hydroxyethyl)-1-piperazine-ethane-sulfonic acid (HEPES) were purchased from Sigma Chemical (St. Louis MO). Draq5 was from Biostatus (Leicestershire, UK), Metamorph image-analysis software was from Molecular Devices (Downington, PA).

**Table 1.**
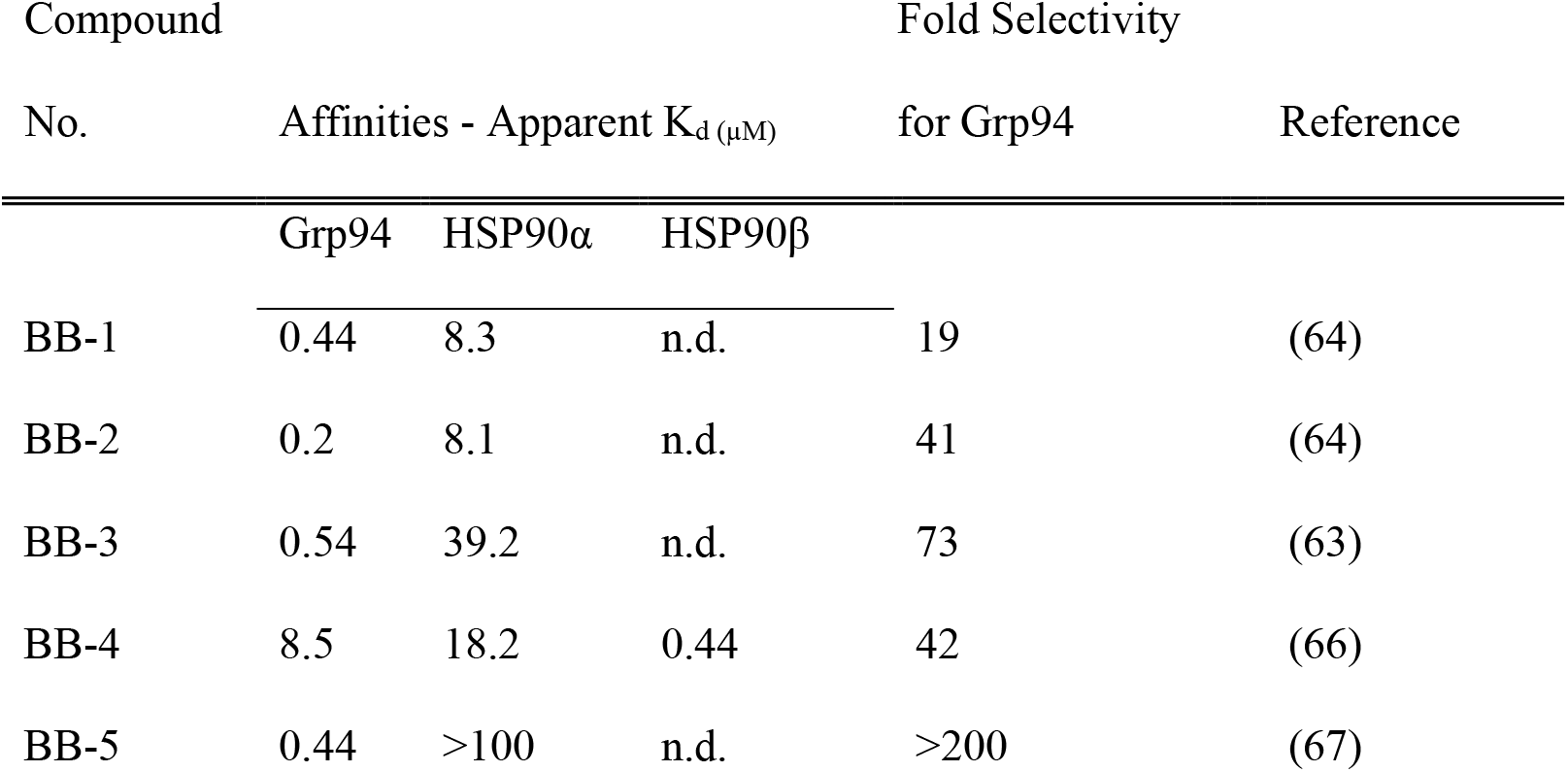
Grp94 Inhibitors

### Cell Culture

#### Human fibroblast cells

Human NPC1 fibroblasts GM18453 (homozygous *NPC1* mutant *I1061T*) were from Coriell Institute, (Camden, NJ). All human skin fibroblasts were maintained in MEM supplemented with 10% FBS. For the inhibitor treatments, cells were maintained in MEM supplemented with 5.5% FBS and 20 mM HEPES.

#### NPC1 Null Cells (U2OS-SRA^shNPC1^)

Stable U2OS-SRA-shNPC1 cells (18) were cultured in McCoy’s 5A medium supplemented with 10% FBS, 50-units/ml penicillin, 50 μg/ml streptomycin, 5 μg/ml puromycin and 1 mg/ml G418.

### Effect of HSP90 inhibitor treatment on human fibroblasts

The dose dependence of six HSP90 inhibitors (17-AAG, AUY922, Ganetespib, AT13387, SNX2112, TAS-116) was determined after 72 h treatment of NPC1 fibroblasts. Human *NPC1^I1061T^* fibroblasts were seeded in 384 well plates at 450 cells/ well in growth medium on day one. After overnight incubation, compounds were added at six different doses diluted in growth medium supplemented with 20 mM HEPES buffer and 5.5% FBS. DMSO was used as a control in each plate for each concentration. After 72 h, the plate was washed with PBS three times, fixed with 1.5% PFA, and stained with 50 μg/ml filipin and nuclear stain Draq5. Measurements were made from four wells for each condition in each experiment, and the experiment was repeated three times. Images were acquired on an ImageXpress^Micro^ automatic fluorescence microscope, at four sites per well and analyzed to obtain the LSO (lysosome like storage organelles) compartment ratio, which is a measure of filipin labeling of stored cholesterol per cell (39). A high LSO ratio is associated with high levels of cholesterol in LE/Ly. Removal of this cholesterol leads to reduction in the LSO ratio. The LSO compartment ratio for each concentration was normalized to corresponding DMSO treated control.

#### Time and Dose Dependence Assay

The dose dependence of two HSP90 inhibitors, 17-AAG and AUY922, was determined as a function of time after 24, 48 and 72 h treatment of *NPC1^I1061T^* fibroblasts (GM18453). To maintain the same density of cells at the final time point, compounds were added chronologically. After overnight incubation, compounds were diluted in growth medium supplemented with 20 mM HEPES buffer and 1% FBS and added in the first plate (for 72 h time point). In the second plate, compounds were added 48 h after seeding the cells and allowed to incubate for 48 h. In the third plate, compounds were added 72 h after seeding the cells and allowed to incubate for 24 h. DMSO was used as a control in each plate for each concentration. All three plates were rinsed, fixed, stained and analyzed as described above.

### Fluorescence Microscopy

An automated ImageXpress^Micro^ imaging system from Molecular Devices equipped with a 300W Xenon-arc lamp from Perkin-Elmer, 10X Plan Fluor 0.3 numerical aperture objective from Nikon, and a Photometrics CoolSnapHQ camera (1,392 x 1,040 pixels) from Roper Scientific was used to acquire images. Filipin images were acquired using 377/50 nm excitation and 447/60 nm emission filters with a 415 dichroic long-pass filter. GFP images were acquired using 472/30 nm excitation and 520/35 nm emission filters with a 669 dichroic long-pass (DCLP) filter. Filter sets assembled in Nikon filter cubes were obtained from Semrock (Rochester, NY).

Images were also acquired using a Leica DMIRB microscope (Leica Mikroscopie und Systeme GmbH, Germany) equipped with a Princeton Instruments (Princeton, NJ) cooled CCD camera driven by Image 1/MetaMorph Imaging System software (Universal Imaging Corporation, PA). Images were acquired using a high-magnification oil immersion objective (63×, 1.4 NA). Filipin images were acquired using an A4 filter cube obtained from Chroma Technology Corp. (Brattleboro, VT) with 360 nm (20-nm bandpass) excitation filter and 465-nm (40-nm bandpass) emission filter.

### Imaging and analysis

A method described previously was used to quantify the cholesterol accumulation in lysosomal storage organelles (LSOs). *NPC1* mutant cells show a bright region of filipin labeling near the cell center, corresponding to the sterol-loaded LSOs. The 384-well plates containing cells were imaged for filipin using a 10X 0.3 NA dry objective on an ImageXpress^Micro^ automatic fluorescence microscope. Four images were acquired from each well. Images were analyzed using MetaXpress image-analysis software. First, all images were corrected for slightly inhomogeneous illumination as described previously (39). A background intensity value was set as the fifth percentile intensity of each image, which corresponds to intensity outside the cells, and this intensity value was subtracted from each pixel in the image. At the plating density used in this study, all fields maintain at least 5% of the imaged area cell-free. We calculated the LSO ratio, which is the ratio of filipin fluorescence intensity in the brightly labeled centers of the cells in the field divided by the total area of the cells. The LSO ratio is determined using two thresholds that are applied to the filipin images. A low threshold is set to include all areas occupied by cells. The outlines of cells using the low threshold are similar to the cell outlines in transmitted light images. A higher threshold is then set to identify regions brightly stained with filipin in cells. The thresholds were chosen for each experiment.

LSO Ratio = Total intensity above high thresholded filipin intensity/ Number of pixels above low thresholded filipin intensity

### Amplex Red Assay for Cholesterol

GM18453 cells were plated in 96 well plates and treated with DMSO or AUY922 at different concentrations for 72 h. For free cholesterol quantification, an Amplex Red assay kit (Abcam Cat# ab65359) was used following the manufacturer’s protocol for lipid extraction and fluorometric measurement.

### NPC1 immuno-localization in NPC1^I1016T^ human fibroblasts

GM18453 cells were treated with 100 nM AUY922 or DMSO solvent control for 72h. Cells were then incubated with 50 μg/ ml Alexa546-LDL in MEM growth medium supplemented with 5.5% FBS and 100 nM AUY922 or DMSO solvent control for 4 h, rinsed with growth medium and chased in MEM growth medium supplemented with 5.5% FBS for 30 minutes. Cells were washed three times with PBS and then fixed with 1.5% PFA in PBS. For immuno-staining cells were permeabilized with 0.5% saponin and 10% goat serum (GS) in PBS for 30 minutes. Cells were incubated with 0.8 μg/ml p-Rab anti-hNPC1 primary antibody for 2 h in the presence of 0.05% saponin and 0.5% GS at room temperature, followed by Alexa488 labeled goat anti-rabbit secondary antibody (1:1000, Life Technologies, Grand Island, NY) for 45 min at room temperature. Finally, cells were washed three times with PBS, and images were acquired using a Leica epifluorescence microscope with a 63X 1.32 NA oil immersion objective and standard FITC and TRITC filters.

### Endoglycosidase H assay

Vehicle control or HSP90 inhibitor-treated NPC1^I1061T^ mutant fibroblasts were lysed with RIPA lysis buffer and endoglycosidase H (EndoH) assay was performed as described previously (40). For EndoH analysis of NPC1 protein, 20 *μ*g of cell lysates were incubated in 4X SDS sample buffer in the presence or absence of EndoH overnight at 37°C. The lysates were resolved on SDS-PAGE followed by immunoblotting with p-Rab anti-hNPC1, p-Rab anti-HSP90, and p-Rab anti-GAPDPH antibodies. The fraction of the intensities of EndoH-sensitive (EndoH^S^) and EndoH-resistant (EndoH^R^) bands for each condition were quantified and plotted.

### Western blotting

Western blot analysis was performed on HSP90 inhibitor treated human NPC1 mutant fibroblasts to measure the expression of NPC1, HSP40, HSP70, and HSP90 using rabbit polyclonal antibodies to NPC1, HSP40, HSP70, and HSP90. Anti-α tubulin was used as a loading control. Secondary antibodies were from Pierce for ECL or from Molecular Devices for Eu label.

### Overexpression of HSP40 and HSP70

For HSP40, NPC1 human fibroblasts were transfected with either eGFP-Vector (pCMV6-AC-GFP Tagged Cloning Vector, Cat #PS100010, Origene Technologies) or eGFP-HSP40 (DNAJB11 (NM_016306) Human Tagged ORF Clone, Cat# RG200216, Origene Technologies) plasmid using an Amaxa human dermal fibroblast kit and T20 protocol. For HSP70, NPC1 human fibroblasts were transfected with either eGFP-Vector or eGFP-HSP70 (Addgene, Cat#15215) plasmid using an Amaxa human dermal fibroblast kit and manufacturer recommended U2OS protocol. After nucleofection, cells were plated in 2 cm poly D-lysine coated cover-slip dishes. Cells were incubated for 72 h, washed with PBS, fixed with 1.5% PFA and stained with 50 μg/ml filipin. Wide-field fluorescence microscope images were acquired using FITC and UV filters and a 10X objective. Images were analyzed to determine LSO ratio values in GFP-labeled cells as described above.

## Results

### Activity of HSP90 inhibitors in lowering lysosomal cholesterol storage in NPC1 mutant cells

We tested the effects of six HSP90 inhibitors on lysosomal cholesterol accumulation in NPC1^I1061T^ mutant human fibroblasts. Cells plated in 384 well plates were treated with 17-AAG (41), AUY922 (41), Ganetespib (42), AT13387 (43–45), SNX2112 (46) or TAS-116 (47) at concentrations ranging from 0.5 nM to 10 μM depending on the reported potency of the compounds. The fibroblasts were treated with inhibitors for 72 h followed by fixation, staining with filipin and Draq5, and fluorescence microscopy analysis. Free (i.e., unesterified) cholesterol levels in perinuclear compartments brightly labeled by filipin in lysosomal storage organelles (LSOs) were measured as described previously (21, 39). All data were normalized to matched, DMSO controls, so a value of one indicates no correction, and values below one indicated reduction of stored free cholesterol. Figure 1A shows the effects of six HSP90 inhibitors. All the inhibitors tested showed dose-dependent effects after a 72 h treatment. AUY922 was the most potent compound, and it partially corrects the NPC1 phenotype below 0.5 nM. The dotted line on the plot indicates DMSO control and the value of one means no correction. The effectiveness of some of the HSP90 inhibitors was reduced at higher concentrations, and we saw a similar reversal with high concentrations of HDACi (21). This may be due to the pleiotropic effects of HSP90 inhibition. The effective concentrations of the HSP90 inhibitors in this assay were generally comparable to values for blocking the growth of cancer cells in culture (43, 44, 46–48), although there was variability of the potency of the compounds in blocking different tumor cells. Detailed statistical analysis was performed using GraphPad PRISM software and the p-values calculated using ANOVA Kruskal-Wallis multiple comparison test are shown in supplementary Figure S1 (A-F). Representative low magnification images of GM18453 cells treated with DMSO or optimal concentrations all HSP90 inhibitors are shown in Figure 1B. Images shown were acquired at 10X magnification on automated high throughput microscope ImageXpress^Micro^. Representative high magnification images acquired using a 63X objective are also shown in Figure 1C. The images in the figure are of NPC1 skin fibroblasts GM18453 treated with DMSO and HSP90 inhibitors at optimal concentrations labeled with filipin. All HSP90 inhibitors tested show substantial reduction in free cholesterol in LSO compared to DMSO treated cells. The reduction in free cholesterol in HSP90 inhibitor treated cells was similar to the reduction we reported previously for cells treated with 10 μM Vorinostat (21).

**Figure 1.**
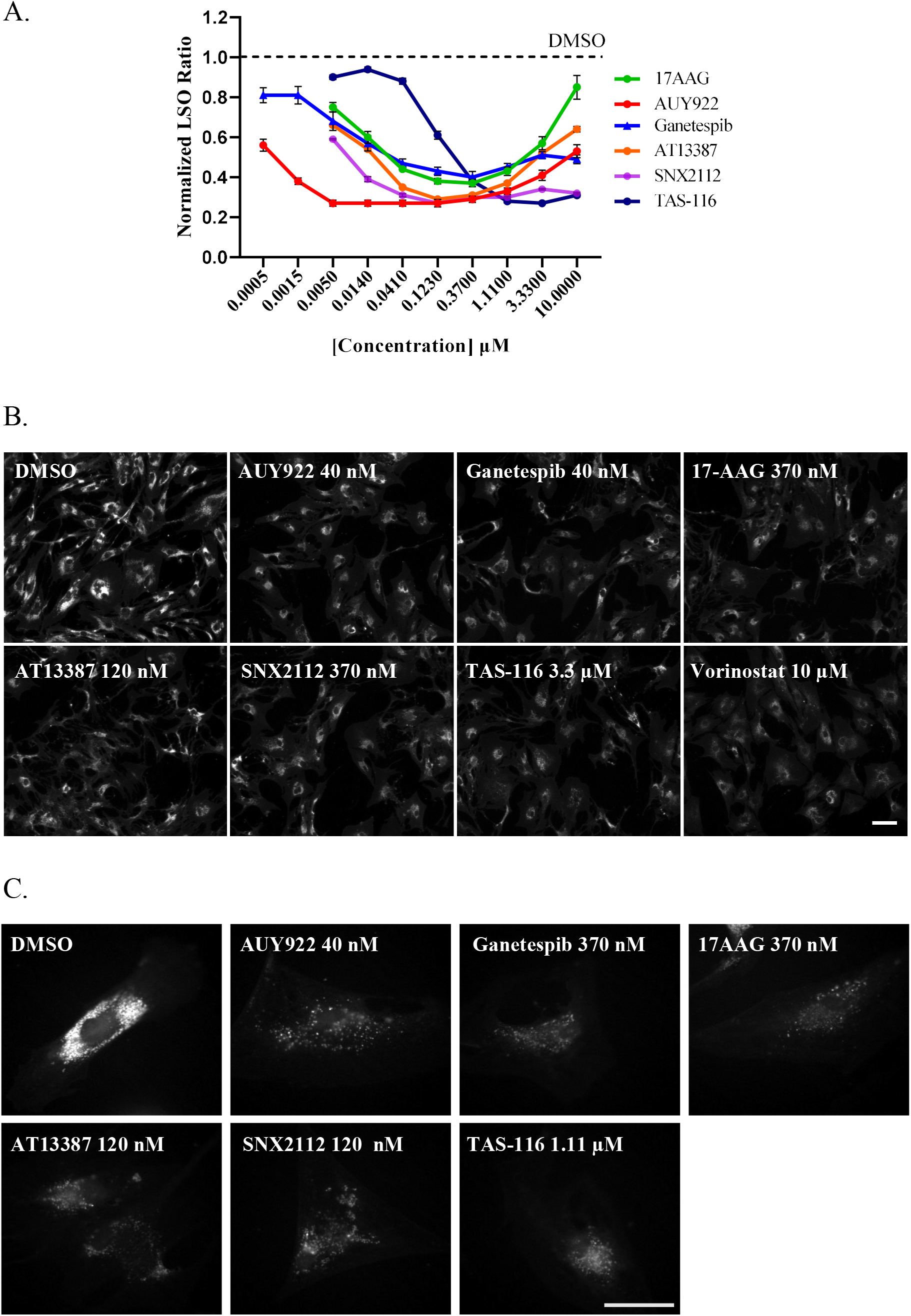
Effect of HSP90 inhibitors on GM18453 NPC1 human fibroblasts: **A.** GM18453 fibroblasts (homozygous I1061T) cells were treated with HSP90 inhibitors 17-AAG, AUY922, Ganetespib, AT13387, SNX2112, and TAS-116 at various doses for 72 h, fixed, and stained with filipin and Draq5. Images were acquired on an ImageXpress ^Micro^ using 10X magnification. Each data point is an average of the LSO values from 48 images. Each image contains about 500-700 cells. Each data point in the plot is from three independent experiments +/- SE. LSO values in DMSO-treated cells were measured in each experiment for normalization and shown as a dotted line. The value of one indicates no correction. **B.** Representative low magnification high throughput filipin images of GM18453 cells treated with DMSO or optimal concentrations of HSP90 inhibitors acquired using 10X objective are shown. Vorinostat treatment was used as positive control. The filipin intensity in perinuclear LSOs of cells treated with HSP90 inhibitors is reduced, demonstrating the clearance of cholesterol. DMSO, AUY922 (40 nM), Ganetespib (40 nM), 17AAG (370 nM), AT13387 (120 nM), SNX2112 (370 nM), TAS-116 (3.33 μM), 17AAG (370 nM), Vorinostat (10 μM), Size bar = 100 μm. **C.** Representative high magnification filipin images of NPC1 GM18453 fibroblast treated with HSP90 inhibitors at optimal concentrations acquired using 63X objective are shown. DMSO, AUY922 (40 nM), Ganetespib (370 nM), 17AAG (0.37 μM), AT13387 (120 nM), SNX2112 (120 nM), TAS-116 (1.11 μM), Size bar = 50 μM

We compared changes in free cholesterol in GM18453 cells using filipin and an Amplex Red assay of cholesterol. Since the Amplex Red assay measures total free cholesterol, we compared this with the total filipin fluorescence per cell. As shown in Figure 2A there is only a slight drop in total cholesterol as the LSOs are cleared of cholesterol, and the filipin and Amplex Red values show very similar effects. The small effect on total cholesterol is in agreement with earlier studies that showed that NPC mutant cells have elevated cholesterol in the LSOs, but elsewhere the cholesterol content is reduced (49). To test whether the observed effect depended on the expression of the mutant NPC1 protein, we treated U2OS-SRA^shNPC1^ (*NPC1^-/-^*) cells with three HSP90 inhibitors at varying concentrations for 72 h. The LSO ratios were measured by filipin labeling as described above. Figure 2B shows that there was no correction of the NPC1 phenotype in these cells, demonstrating that the effect observed is specific for cells carrying a mutant NPC1 protein and not an NPC1-independent effect similar to HPCD treatment (50).

**Figure 2.**
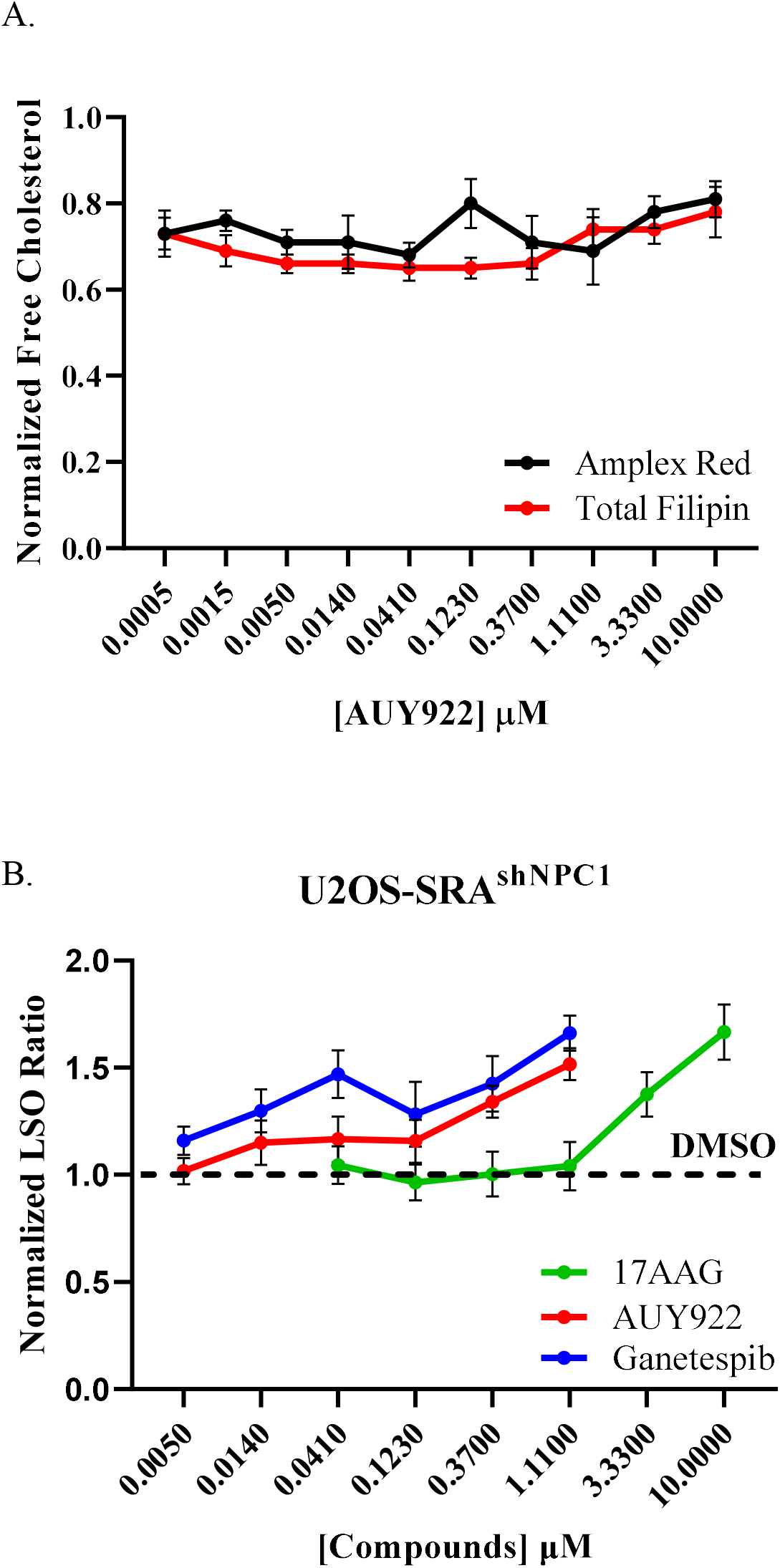
**A.** *Total cholesterol estimation.* A Plot showing total cholesterol concentration in AUY922 treated GM18453 cells as measured by Amplex Red biochemical method (black) and Average filipin per cell (red). Error bar = SE. **B.** *Treatment of HSP90 inhibitors on U2OS^shNPC1^ cells*. U2OS-SRA-shNPC1 cells were treated with HSP90 inhibitors at varying doses ranging from 5 nM to 10 mM for 72 h, fixed and stained with Filipin and Draq5. Images were acquired on an ImageXpress ^Micro^ using a 10X objective. Each data point is an average of 48 images. Each image containing ~500-700 cells. Each data point in the plot is from three independent experiments. Error bar = +/- SE.

### Time Course of HSP90 inhibitors on NPC1^I1061T^ fibroblasts

We measured the time course for effects of HSP90 inhibitors on NPC1^I1061T^ mutant human fibroblasts. Figure 3 shows the effect of 17-AAG and AUY922 at various doses after 1, 2, or 3 days. Neither of the inhibitors tested had a large effect at 24 h, but both showed dose-dependent effects at 48 h and similar or somewhat greater effects after 72 h treatment. This time course is consistent with the need to synthesize and transport new NPC1 proteins to LE/Ly in response to the drug treatment as has been proposed for HDACi treatment (11, 18, 21). The t1/2 of wild type NPC1 wild type protein that has passed beyond the ER is 1-2 days (11), and a similar lifetime was observed for NPC1^I1061T^ in human fibroblasts treated with Vorinostat (18).

**Figure 3.**
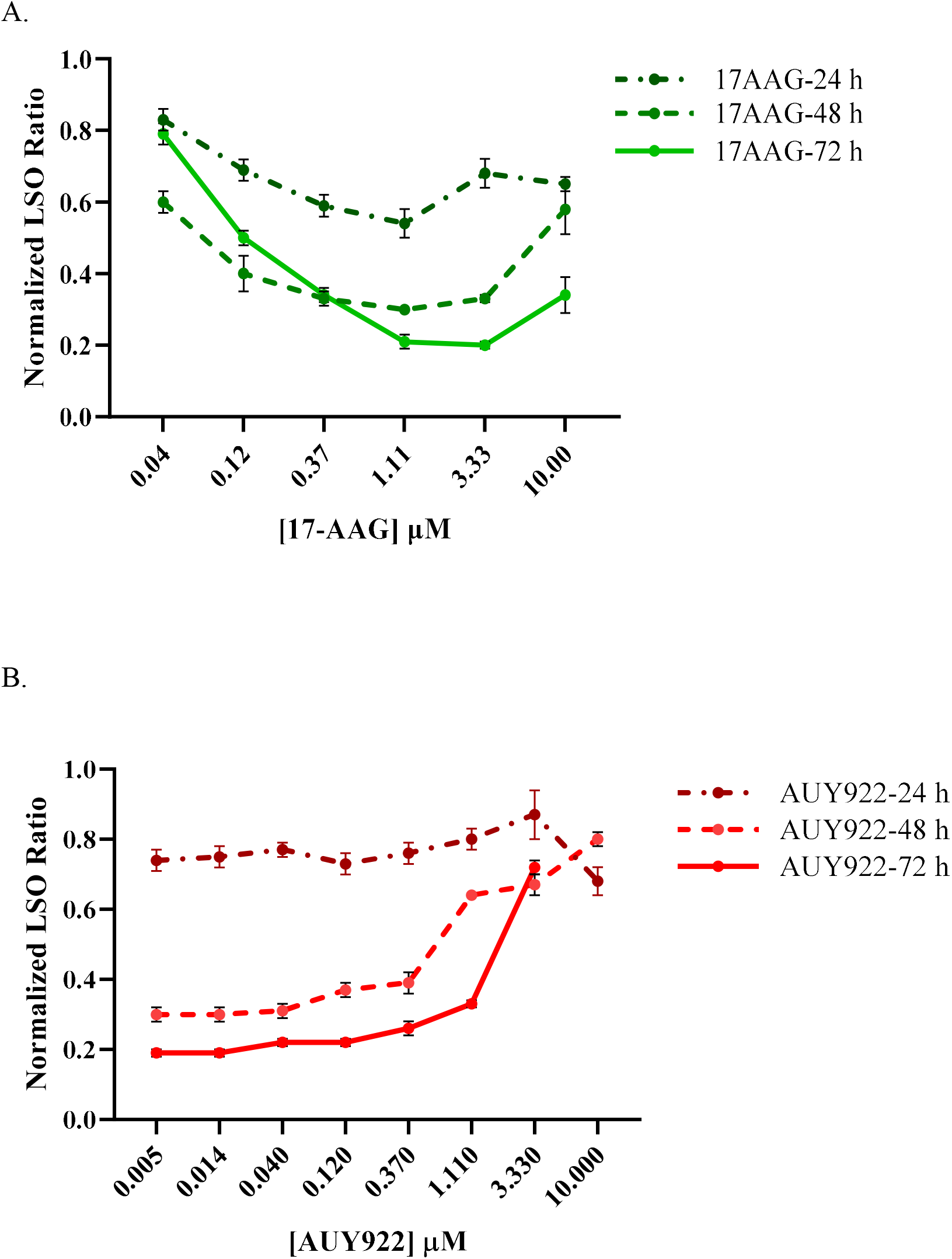
Time course of treatment with HSP90 inhibitors: GM18453 fibroblasts were treated with **A.** 17-AAG or **B.** AUY922 at varying doses for 24, 48 and 72 h, fixed, and stained with filipin and Draq5. Images were acquired on an ImageXpress ^Micro^ using a 10X objective. Each data point is an average of 500-700 cells per image and 48 images. Each data point in the plot is from three independent experiments. Error bar = +/- SE.

### AUY922 treatment rescues the trafficking of mutant NPC1 protein in NPC1^I1061T^ fibroblasts

In cultured fibroblasts approximately half of the wild type NPC1 protein is degraded rapidly, and for many mutants most of the synthesized protein is degraded with very little reaching the LE/Ly (11, 18). To measure export of NPC1^I1061T^ out of the ER, we measured acquisition of resistance to endoglycosidase H (EndoH), which occurs after proteins are exported from the ER. We measured the effect of AUY922 on acquisition of EndoH resistance (EndoH^R^). The cell lysates from DMSO and AUY922 treated NPC1I^1061T^ fibroblasts were incubated with endo H to cleave immature high-mannose *N*-linked glycans found in the ER, resulting in an increase in migration using SDS-PAGE (EndoH^S^). In immunoblots, the band intensities for EndoH^R^ and EndoH^S^ bands were measured (Figure 4 A,B). Treatment of NPC1I^1061T^ fibroblasts with AUY922 substantially increases exit from the ER as shown by the decrease in EndoH^S^ fraction (clear bars) after AUY922 treatment in a dose dependent manner.

**Figure 4.**
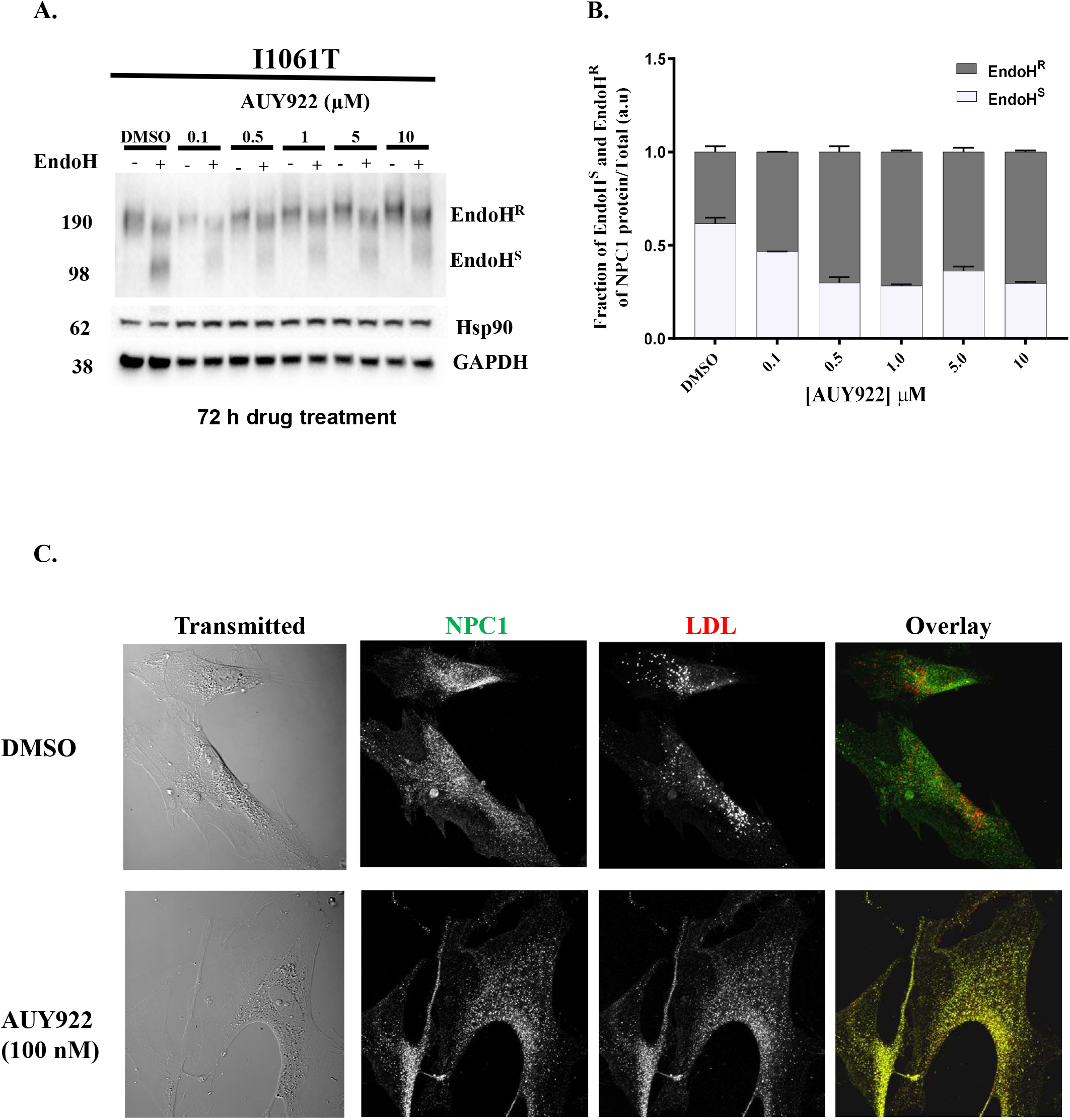
Localization and stability of NPC1 protein after HSP90 inhibitor treatment: **A.** GM18453 cells were treated with AUY-922 at varying doses for 72 h, after which cells were lysed with RIPA lysis buffer. For EndoH analysis of NPC1 protein, 20 *μ*g of cell lysates were treated in the presence or absence of EndoH overnight at 37°C. The lysates were resolved on SDS-PAGE followed by immunoblotting with p-Rab anti-hNPC1, p-Rab anti-HSP90 and p-Rab anti-GAPDH antibodies. **B.** The fraction of the intensities of EndoH^S^ and EndoH^R^ bands for each condition. **C.** GM18453 fibroblasts were treated with 100 nM AUY-922 or solvent control DMSO for 72 h. The cells were incubated with Alexa-546 LDL for 3.5 h followed by a 0.5 h chase to deliver labeled LDL to LE/Ly. The fibroblasts were fixed, and immuno-stained with primary p-Rab-anti hNPC1 antibody followed by goat-anti-Rab-A488 labeled secondary antibody and detected by immunofluorescence. Images were acquired by confocal microscopy. Representative transmitted light and sum projected images of fluorescently labeled NPC1 and LDL are shown. Size bar = 25 μm.

To determine whether treatment with an HSP90 inhibitor improved trafficking of NPC1^I1061T^ protein to LE/Ly, we treated GM18453 NPC1^I1061T^ cells with 100 nM AUY922 or solvent control DMSO for 72 h. The cells were then incubated with Alexa546 LDL for 3.5 h followed by a 0.5 h chase to deliver labeled LDL to LE/Ly (51–53). The fibroblasts were fixed, and NPC1 protein was detected by immunofluorescence (Figure 4C). Representative sum projected images are shown. The mutant NPC1 protein does not co-localize with Alexa546 LDL in LE/Ly in the DMSO-treated cells (*i.e*., red and green puncta are not co-localized). After 72 h treatment with 100 nM AUY922, a large fraction of the mutant NPC1 protein localizes in yellow puncta in the overlay, showing that NPC1 and LDL are in the same organelles. Additional representative images are also shown in Figure S2. This indicates that treatment with AUY922 leads to delivery of the mutant NPC1 to the organelles in which LDL is releasing cholesterol, and this NPC1^I1061T^ effectively aids in transporting cholesterol out of these organelles.

### Effect of HSP90 inhibitor treatment on expression of other proteins

Chaperone-mediated protein folding involves synchronized interplay of multiple chaperones and co-chaperones, so inhibiting one protein results in perturbation/modulation of other chaperones and co-chaperones in the pathway. It has been reported that inhibiting HSP90 results in upregulation of HSP70 in cells (54, 55). HSP90 inhibitors do not generally change the expression of the HSP90 protein, but they inhibit its activity by binding at the N-terminal catalytic ATP binding site (48, 56–58). To further understand the mechanism by which HSP90 inhibitors reverse the NPC1 mutant phenotype, we measured the expression of HSP90, HSP70, and HSP40 as well as the NPC1 protein.

NPC1^I1061T^ human fibroblasts were treated with either 17-AAG (1μM), AUY922 (40 nM), Ganetespib (100 nM), or DMSO (vehicle control). After 72 h treatment with compounds or solvent control, cells were lysed, followed by immunoblot analysis. Figure 5A shows representative bands from one experiment. Average band intensities for three independent experiments were quantified and normalized to loading control (tubulin) and compared to the parallel DMSO-treated band (Figure. 5B). The dots shown on each bar represents values from independent experiments and the data plotted are +/- SE. The results indicate that HSP90 inhibitor treatment results in significant increases in expression of NPC1, HSP70 and HSP40, but there is no significant change in HSP90 expression. The increase in HSP70 and HSP40 expression after HSP90 inhibitor treatment is in agreement with previously published reports (12, 54, 55, 59, 60).

**Figure 5.**
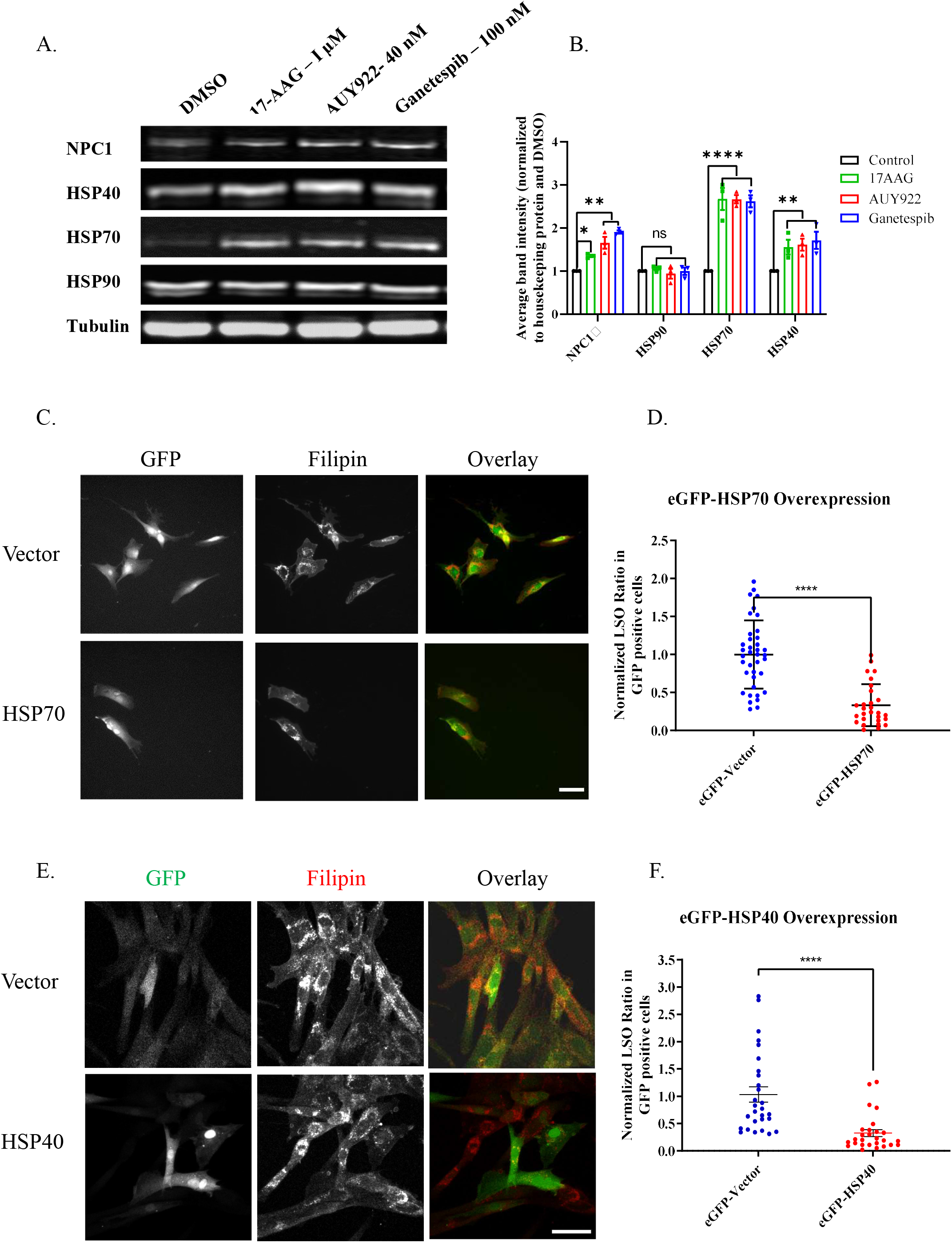
Effect of HSP90 inhibitors on expression of other chaperones: GM18453 fibroblasts were treated with 17-AAG (1μM), AUY922 (40nM), or Ganetespib (100 nM). As a control, cells were treated with DMSO. After 72 h treatment with compounds or solvent control, cells were lysed and immunoblotted for various proteins. After resolving bands on 4-12% Bis-Tris gel, proteins were transferred to a PVDF membrane and probed with p-Rab anti-hNPC1 antibody, p-Rab anti HSP70, p-Rab anti HSP40 and p-Rab anti HSP90. Tubulin (mouse monoclonal) was used as a loading control. **A.** Lane 1: DMSO; Lane 2: 1μM 17-AAG; Lane 3: 40 nM AUY922; and Lane 4: 100 nM Ganetespib. **B.** Plot showing quantification of immunoblot bands normalized to tubulin and DMSO treated fibroblasts. Dotted indicates DMSO control. P values for NPC1, HSP90, HSP70 and HSP40 were 17-AAG: 0.001, ns, 0.008, 0.05; AUY922: 0.03, ns, 0.0002, 0.03 and Ganetespib: 0.0001, ns, 0.0009,0.05 respectively. **C.** GM18453 cells were transiently transfected with eGFP-Vector or eGFP-HSP70 and incubated for 72h. Cells were washed with PBS, fixed with 1.5% PFA and stained with 50 μg/ml filipin. Images were acquired using GFP and A4 filters using a 20X dry objective. Size bar = 25 μm. **D.** LSO filipin was measured in GFP positive cells and plotted. Dot plots showing data from three independent experiment +/- SE, Mann-Whitney statistical test was performed, p<0.0001. **E.** GM18453 cells were transiently transfected with eGFP-Vector or eGFP-HSP40 and incubated for 72 h. Cells were washed with PBS, fixed with 1.5% PFA and stained with 50 μg/ml filipin. Images were acquired using GFP and A4 filters using a 20X dry objective. Size bar = 10 μm. **F.** LSO filipin was measured in GFP positive cells and plotted. Dot plots showing data from three independent experiment +/- SE, Mann-Whitney statistical test was performed, p<0.0001.

### HSP70 (HSPA1A) Overexpression

In order, to determine whether increased HSP70 expression plays a role in correction of the NPC1^I1061T^ mutant phenotype, we overexpressed HSP70 in NPC1^I1061T^ human fibroblasts. We transiently transfected eGFP-HSP70 in GM18453 cells. Representative images for eGFP-Vector transfected and eGFP-HSP70 transfected cell are shown in Figure 5C. Figure 5D shows the plot of LSO Ratio in GFP-positive cells that did or did not overexpress HSP70. The data plotted are from two independent experiments and 40 images. These results indicate that the increased expression of HSP70 aids in rescuing the *NPC1^I1061T^* mutant phenotype.

### HSP40 (DNAJB11) Overexpression

HSP40 is a co-chaperone of HSP70, so we tested the role of HSP40 expression in the correction of NPC1^I1061T^ mutant phenotype. We overexpressed HSP40 in NPC1^I1061T^ human fibroblasts by transiently transfecting eGFP-HSP40 in GM18453 cells. Representative images for eGFP-Vector transfected and eGFP-HSP40 transfected cells are shown in Figure 5E. Figure 5F shows a plot of the LSO Ratio in GFP-positive cells that did or did not overexpress HSP40. The data are from three independent experiments and 24-30 images. These results indicate that the increased expression of HSP40 also results in rescuing the *NPC1^I1061T^* mutant phenotype.

### Grp94 Inhibitors

The HSP90 family of genes includes HSP90A (cytosolic), HSP90B (ER) and TRAP (mitochondrial) proteins. The inhibitors that we described so far are pan-inhibitors or just inhibit HSP90A (TAS-116) (61). The TAS-116 results indicated that inhibition of the cytosolic form of HSP90 alone was effective. In order to examine another HSP90 isoform we tested inhibitors selective for the ER resident HSP90B1 class of chaperones, GRP94 (62–67). GM18453 fibroblasts were treated with five different GRP94 inhibitors (Table 1) for 72h, and the LSO Ratio was measured (Figure 6). The GRP94-selective compounds did not rescue the NPC1^I1061T^ phenotype. These data indicate specificity for cytosolic HSP90A inhibition in rescuing NPC1 phenotype.

**Figure 6.**
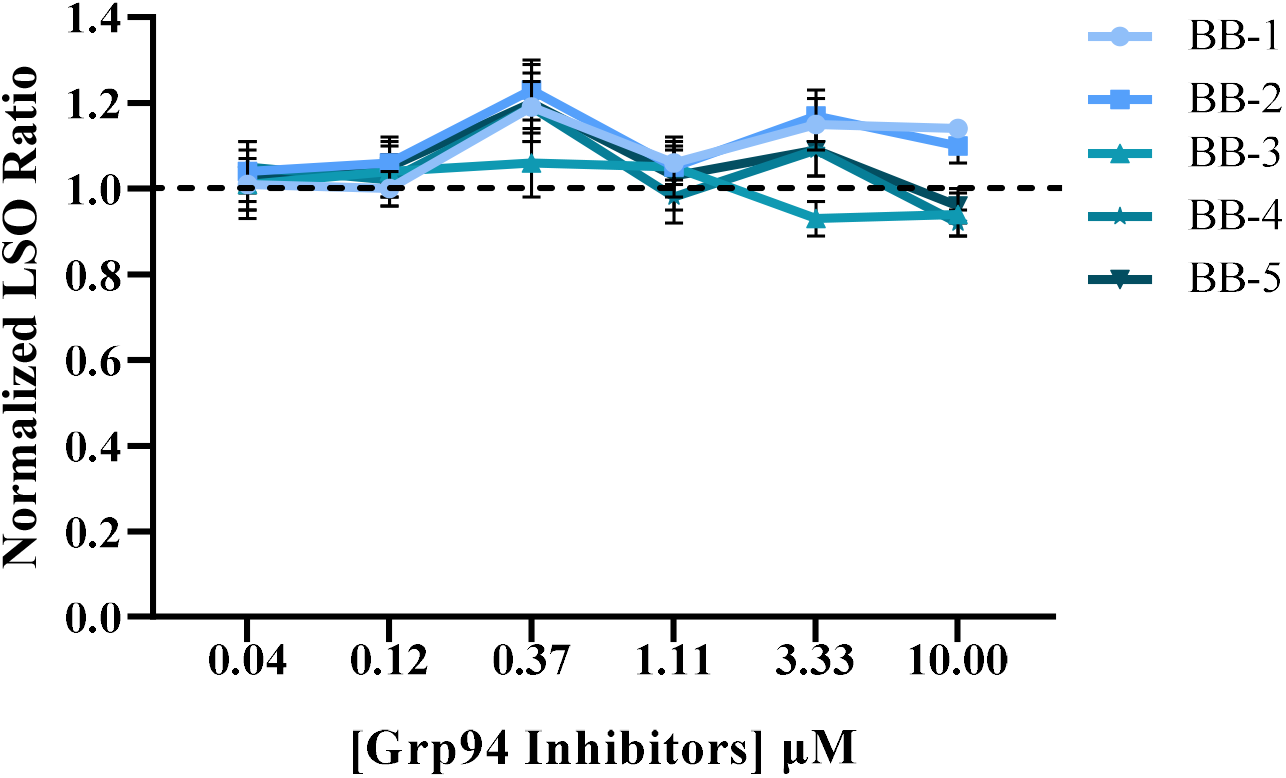
Effect of GRP94 Inhibitors on NPC1 cells: GM18453 fibroblasts were treated with GRP94 inhibitors BB1, BB2, BB3, BB4 and BB5 at varying doses for 72 h, fixed, and stained with filipin and Draq5. Images were acquired on ImageXpress ^Micro^ using a 10X objective. Each data point is an average from three experiment and 72 images total, each image contained ~300-500 cells. Error bar = +/- SE.

### HSP70 Activators

It has been reported that, arimoclomol, reduces the size of lysosomes in NPC1-null fibroblasts, and it reduced the ataxia and improved behavioral symptoms in an *Npc1^-/-^* mouse model (34). The compound arimoclomol has been described as an HSP70 co-inducer, but we could find no peer reviewed publications supporting this. The efficacy and safety of arimoclomol is being tested on NPC1 patients in a clinical trial (68). We tested arimoclomol on GM18453 human NPC1 fibroblasts at concentrations ranging from 1.6 μM to 500 μM for three days or five days (Figures 7A). We did not observe correction of the cholesterol storage in NPC1^I1061T^ human fibroblasts. At higher concentrations and longer exposure, the phenotype worsened as shown by increases in the LSO Ratio. In order to determine whether arimoclomol upregulates HSP70 protein expression in our cell culture model, we performed Western blot analysis on GM18453 cells treated with arimoclomol (50 and 400 μM) for three and five days (Figure 7B) and plotted the results from three independent experiments (Figure 7C). The results indicate that arimoclomol treatment by itself does not upregulate HSP70 expression in GM18453 fibroblasts.

**Figure 7.**
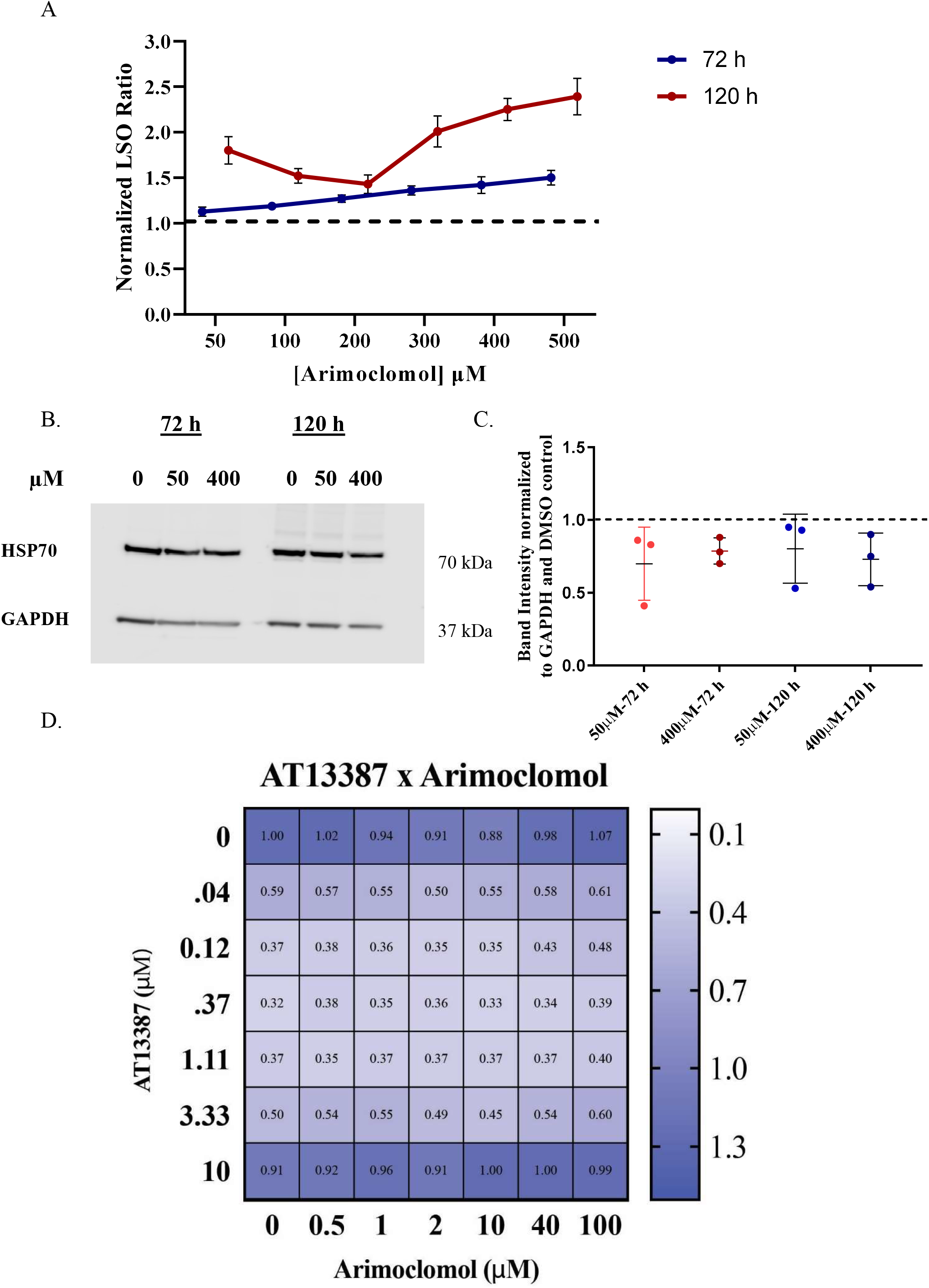
**A**. *Effect of HSP70 co-inducer arimoclomol on NPC1 human fibroblasts*: GM18453 fibroblasts were treated with HSP70 activator arimoclomol for 72 h or 120 h at doses ranging from 50 μM to 500 μM, fixed, and stained with filipin and Draq5. Images were acquired on ImageXpress ^Micro^ using a 10X objective. Each data point is an average of 48 images. Each image containing ~500-700 cells. Each data point in the plot is from three independent experiments +/- SE. **B.** *Western blot of arimoclomol treated NPC1 human fibroblast*: GM18453 fibroblasts were treated with HSP70 activator arimoclomol for 72 h or 120 h at 50 μM to 400 μM. Cells were lysed and Western blot was run using protocol described in method section. Membrane was probed with p-Rab-anti-HSP70 and p-Rab-anti-GAPDH antibodies. Bands were visualized using ECL on Licor Fc system and quantified. **C**. Data in plot are from three independent experiments. There was no significant difference in HSP70 expression compared to DMSO treated cells. Error bars = SE. **C.** Plot showing quantification of immunoblot bands from DMSO or 50 and 400 μM arimoclomol treated NPC1 fibroblasts for 72 h and 120 h. Data were normalized to tubulin and DMSO treated fibroblasts. Dotted indicates DMSO control. No statistically significant difference was observed. **D.** *Cotreatment of AT13387 and arimoclomol on GM18453 NPC1 fibroblasts*. GM18453 fibroblasts were plated in 384-well plates and co-incubated with arimoclomol and HSP90 inhibitor AT13387 in a dose-dependent manner for 72 h. Heatmap value represents the average LSO of three independent experiments with four replicate per wells per plate, each image had 100-300 cells, normalized to DMSO-treated cells. LSO values less than one represent a decrease in cholesterol storage, and values greater than one identify an increase in cholesterol storage.

### Cotreatment of arimoclomol with AT13387 and HSF1a

Since treatment with HSP90 inhibitors increased HSP70 expression, we tested whether arimoclomol in combination with one of the HSP90 inhibitors would show synergistic effects. NPC1 human fibroblasts GM18453 were treated with compounds one day after plating. Cells were treated with arimoclomol at 0, 0.5, 1, 2, 10, 40 and 100 μM in combination with AT13387 (Onalespib) at 0, 0.04, 0.12, 0.37, 1.11, 3.33 and 10 μM and compared against DMSO-treated controls. The range of drug dosages were made to contain a large gradient of concentrations in screening media as described previously (39). Figure 7D shows the heat map of the combination treatments. The treatment with AT13387 did show correction of NPC1 phenotype as expected, but there was no statistically significant synergistic effect from including arimoclomol.

Heat shock transcription factor 1 (HSF1), is the master regulator of Hsp gene transcription (69). arimoclomol is described as a co-activator that prolongs the binding of activated HSF1 to heat shock elements (HSEs) in the promoter regions of inducible Hsp genes (70). A study of beneficiary effects of arimoclomol in NPC1 disease reported that NPC1 fibroblasts subjected to heat shock exhibited activation of HSF1 (34). Hence, we tested the effect of arimoclomol in combination with HSF1 activating compound HSF1a (71, 72) and measured the cholesterol level using the LSO filipin assay. As shown in Figure S3, we did not see a beneficial effect of HSF1a with or without arimoclomol.

## Discussion

We report here that treatment of NPC1 mutant human fibroblasts with several HSP90 inhibitors at nanomolar concentrations leads to clearance of the accumulated cholesterol in the LE/Ly of human homozygous NPC1^I1061T^ fibroblasts in a dose-dependent manner. Based on the ineffectiveness of these inhibitors on NPC1^-/-^ cells, we infer that the clearance is due to a direct impact on the NPC1 protein and not a general bypass of the NPC1-dependent pathways as is seen in cells treated with cholesterol-chelating cyclodextrins (50).

The beneficial effect of HSP90 inhibitors is associated with improved trafficking of the NPC1 protein out of the ER and delivery to LE/Ly. In untreated cells, the majority of NPC1^I1061T^ protein is EndoH sensitive, indicating failure to exit the ER (29). After the treatment with AUY-922 for three days the majority of the NPC1^I1061T^ has exited the ER and shows good co-localization with internalized LDL in LE/Ly. In order, to start understanding how inhibition of HSP90 might improve folding of NPC1^I1061T^ and its exit from the ER, we examined he effects of HSP90 inhibitors on expression of other chaperone proteins. As reported previously, HSP70 and HSP40 levels were increased when HSP90 was inhibited (32, 33), and this was accompanied by an increase in NPC1 protein in the cells. Transient overexpression of HSP70 or HSP40 also reduced the cholesterol storage significantly.

A recent study examined the effects of modulating the HSP70 chaperone/co-chaperone complex on a large number of *NPC1* variants (14). That study focused on the effects of an allosteric inhibitor, JG98, that alters the interactions of HSP70 with co-chaperones, especially the BAG family of co-chaperones (73, 74). As with the study on HDAC inhibitors (18), it was found that JG98 corrected the ER retention and degradation of many *NPC1* variants (14). This study examined SREBP2 activity, which is affected by the release of cholesterol from LSOs, and the results indicate that JG98 was facilitating release of cholesterol from LSOs in *NPC1^I1061T^* cells. Overall, our results corroborate these findings on the importance of HSP70 and other chaperones and extend them by directly demonstrating trafficking of NPC1^I1061T^ to late endosomes and lysosomes containing LDL and showing reduction of filipin-labeled cholesterol in these organelles. Much additional work will be required to develop optimal strategies to target these chaperone networks to treat NPC1 patients.

Since NPC1 protein is folded initially in the ER, we tested inhibitors of the ER-specific member of the HSP90 family, GRP94 (63–65, 67). The GRP94-selective inhibitors were not effective in rescuing the cholesterol storage defect in NPC1^I1061T^ cells. This is consistent with a downstream effect of HSP90 inhibitors, such as increasing cytosolic HSP70 expression, rather than a direct effect of an HSP90 protein on the NPC1 protein in the ER. Our data showing rescue of NPC1 phenotype with HSP40 overexpression is in agreement with published reports. It has been shown that treatment of HEK293 cells with 17AAG increases the expression of HSP70 mRNA, and the effect is enhanced by a mild heat stress (75). It will be valuable in the future to explore treatments that enhance HSP70 activity (and other chaperones) for the treatment of NPC1 disease.

We tested arimoclomol treatment on disease relevant NPC1^I1061T^ cells, which was previously reported to enhance HSP70 as well as HSF1 activation in NPC^-/-^ cells (34). However, we did not observe any reduction in cholesterol storage in NPC1^I1061T^ cells at concentrations up to 0.5 mM for three or five days. Nor did we see any increase in HSP70 expression in NPC1^I1061T^ cells after arimoclomol treatment (Figure 5B, C). A related compound, bimoclomol, has been reported to enhance stress responses but it requires a mild independent activation of stress for this effect (76). It is possible that this type of stress was not present in our cells. Based on the reports of increased HSF1 activation upon arimoclomol treatment and a study reporting combination treatment of arimoclomol with celastrol – an HSF1 activating compound (34, 70), we treated NPC1^I1061T^ cells with a combination of arimoclomol and an HSF1-activating compound (HSF1a) (71, 72). This did not yield any additional correction of the NPC1 phenotype. In contrast when we combined arimoclomol with AT13387 compound, which has shown to correct the NPC1 phenotype (Figure 1), we did observe the expected correction from AT13387, but there was no synergistic enhancement by arimoclomol combination treatment (Figure 7D). Taken together our data suggests that the therapeutic effect of arimoclomol treatment on NPC1^-/-^ cells might be due to off-target effects since arimoclomol has been reported to inhibit several key regulatory enzymes at micromolar concentrations (77).

This result suggests that further exploration of the mechanism of action of arimoclomol is warranted especially since the pharmaceutical form of arimoclomol is currently undergoing clinical trials (68). There are numerous mutations reported in the human *NPC1* gene that are associated with NPC1 disease (2, 3, 5, 7). The most common mutant protein, NPC1^I1061T^, exits the ER inefficiently and is mostly degraded (18). We have previously shown that several other NPC1 missense mutations could function if they could pass through the ER quality control processing. In this study, we only tested NPC1^I1061T^, but we found that the results were very similar to HDACi treatment in that we observed increased protein stability and proper localization of the mutant protein. In conclusion, we report here the therapeutic potential of pan HSP90 inhibitors or HSP90A-selective inhibitors for the treatment of NPC1 disease. Efforts are under way to test these compounds in a mouse model. We note that although TAS-116 is not as potent as other HSP90 inhibitors tested, it is selective for HSP90α and HSP90β (61) in cytosol/nucleus and does not target GRP94 in ER and TRAP1 in mitochondria. In addition, unlike the other HSP90 inhibitors, TAS-116 is an oral drug that is currently in a Phase 3 clinical trial of gastrointestinal stromal cancer (GIST) (78, 79). The promising effect of HSP90 inhibition and HSP70 overexpression provides a basis for further investigations of protein chaperones in the treatment of NPC1 disease.

## Data availability statement

Concept and design NHP, FRM. Acquisition, analysis, and interpretation of data NHP, KS additional experiments SZS AC, AM and KG, Reagents support PH, OW, BB, LN, DSO. Drafting of manuscript NHP, FRM. Critical revision of the manuscript for important intellectual content all authors

## Acknowledgements

This work was supported in part by grants from the NIH, R01-NS092653 (DSO, FRM, PH, OW) and R01-CA213566 (BSJB), by grants from the Ara Parseghian Medical Research Foundation (DSO, FRM, PH, OW), by a grant from MDBR-UPenn Orphan Disease Center (NHP), by a grant from the Harrington Discovery Institute (DSO), and by grants from Dana’s Angels Research Trust (DSO, FRM).

## Abbreviations

LSO: Lysosomal Storage Organelle
NPC: Niemann-Pick type C

**Footnotes to text (if any) - None**

## Supplementary Material

### Methods

#### Statistical Analysis of HSP90 inhibitors dose response plot

The data for HSP90 inhibitor treated GM18453 NPC1 human skin fibroblasts were analyzed for statistical significance using ANOVA Kruskal-Wallis multiple comparison test in Graphpad PRISM software. The p-values compared to lowest concentration in each treatment are shown in Figure S1

#### Co-treatment of Arimoclomol (ACM)l with HSF1 Activator (HSF1a)

Mutant human fibroblasts (GM18453) were purchased from the Coriell Cell Repositories (Camden, NJ) and were grown in Eagle’s MEM with Earle’s salts and 10% FBS growth media. For screening purposes, growth medium supplemented with 5% FBS was used. We seeded GM18453 cells (450 cells/well in 30 μL) in Corning 384-well black polystyrene flat, clear-bottomed tissue culture-treated plates.

Cells were treated with either ACM alone or in combination with HSF1a in a dose-dependent manner and compared against DMSO-treated controls. The range of drug dosages were made to contain a large gradient of concentrations in screening media as described previously (Pipalia, Huang et al. 2006). Briefly, 15 μL of 4x concentration of ACM was combined with either 15 μL of 4x concentration of HSF1a and added to appropriate wells to yield 60 μL of 1x concentration of each drug. In control wells corresponding concentrations of DMSO was used.

All plates were incubated with compounds for 72 h at 37°C. Plates were then washed three times with PBS, pH 7.4, using a Bio-Tek Elx405 plate washer (Bio-Tek Instruments, Inc., Winooski, VT). For each wash cycle, 70 μl of PBS was dispensed followed by aspiration with a residual volume of 16 μl/well. Cells were then fixed with 1.5% PFA in PBS for 20 min at room temperature, followed by three more washes with PBS. To the fixed cells, filipin was added at a final concentration of 50 μg/ml in PBS for 45 min at room temperature. Cells were finally washed three times with PBS followed by addition of 2 μM Draq5. Images were acquired three hours after labeling

Measurements were made from four wells for each condition in each experiment, and the experiment was repeated three times. Images were acquired using a 10X dry objective on an ImageXpress^Micro^ fluorescence microscope at two sites per well and analyzed to obtain the LSO compartment ratio. All data were normalized to DMSO treated values.

**Figure. S1.**
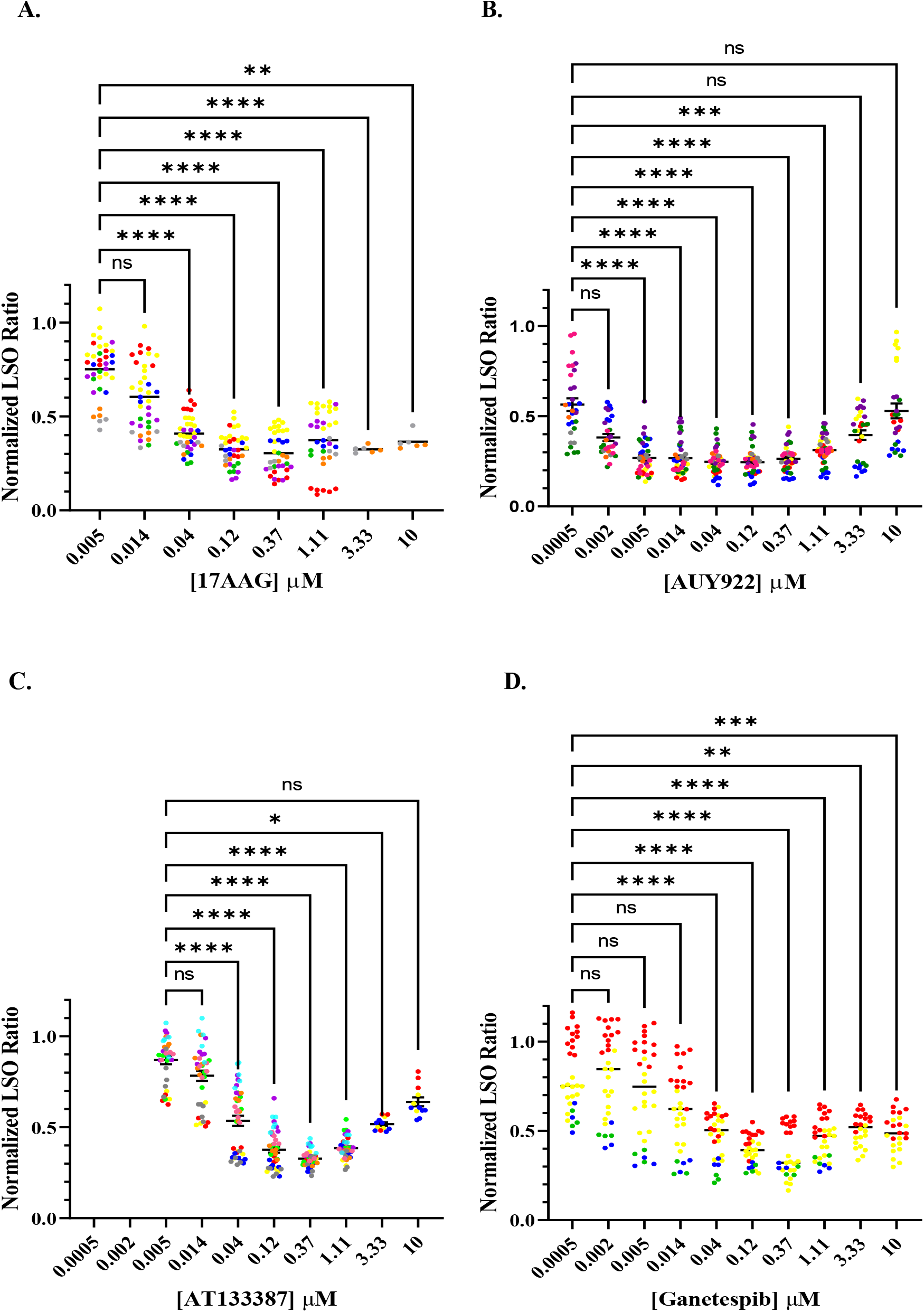
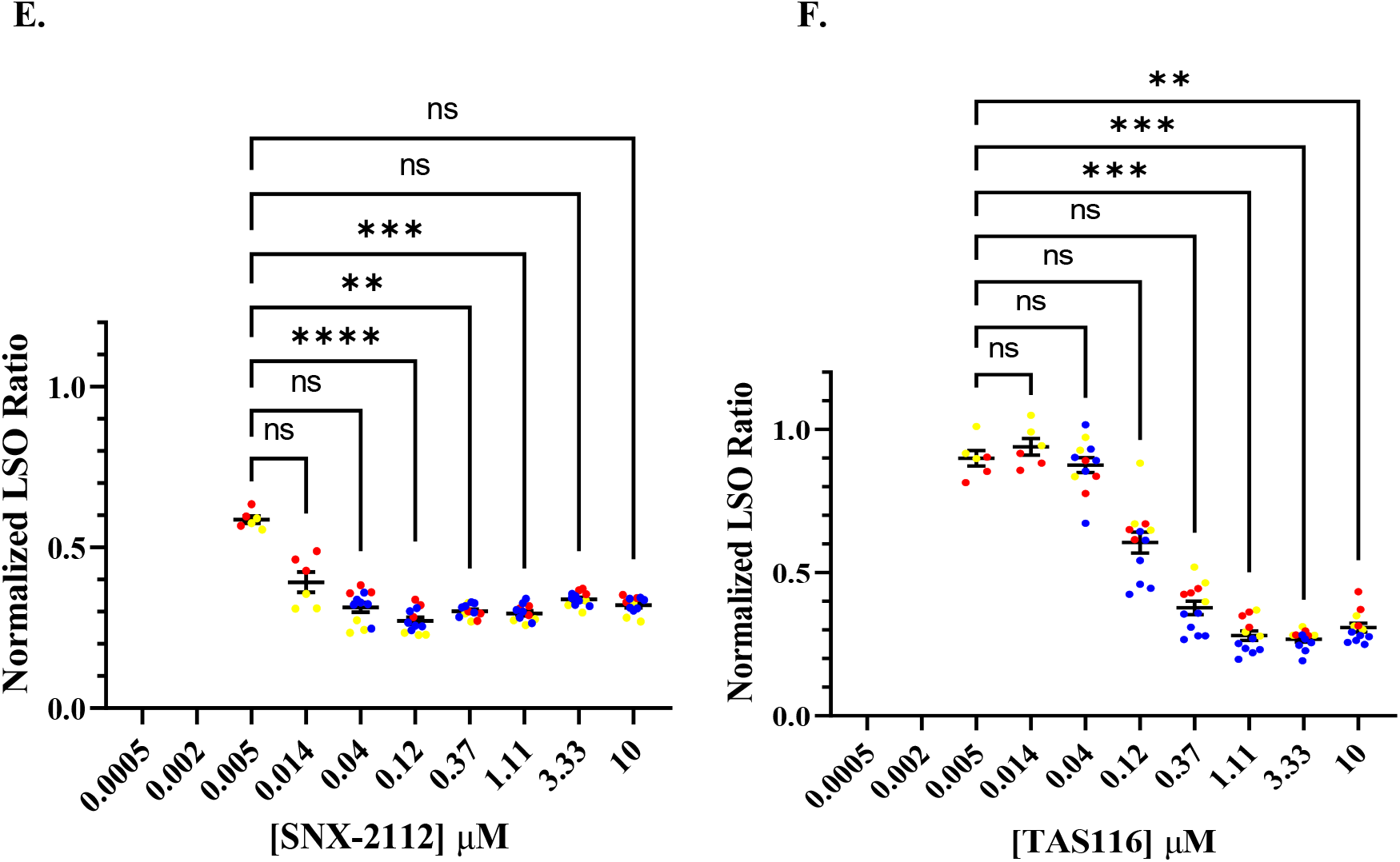
Dot Plot showing statistical significance of dose dependance of various HSP90 inhibitors using ANOVA Kruskal-Wallis multiple comparison test **A-F**. Plots showing statistical significance in dose response curve using ANOVA Kruskal-Wallis multiple comparison test in GraphPad PRISM. **A.** 17AAG, **B**. AUY922, **C.** Ganetespib, **D.** AT13387, **E.** SNX-2112, **F.** TAS116

**Figure S2.**
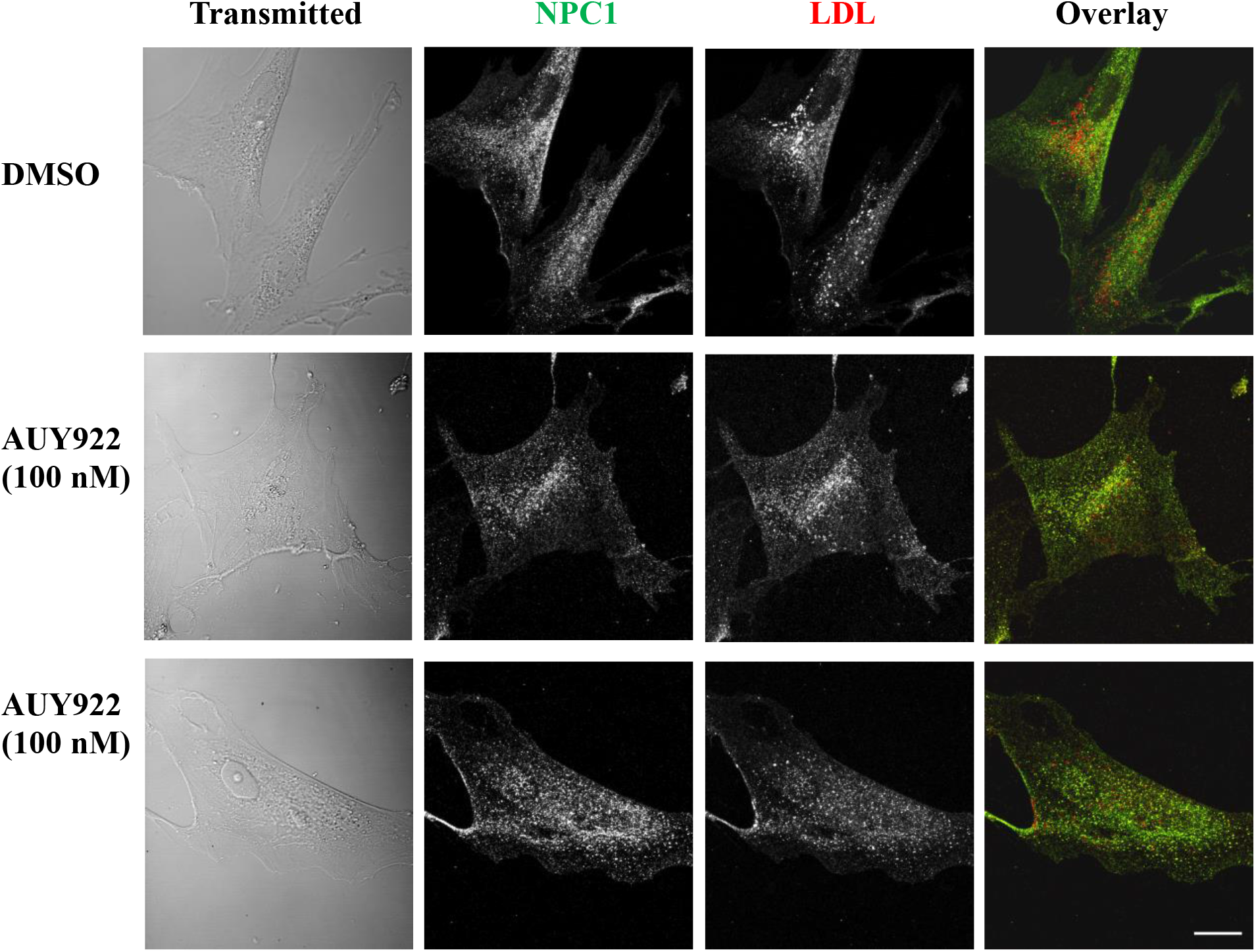
GM18453 fibroblasts were treated with 100 nM AUY-922 or solvent control DMSO for 72 h. The cells were incubated with Alexa546 LDL for 3.5 h followed by a 0.5 h chase to deliver labeled LDL to LE/Ly. The fibroblasts were fixed, and NPC1 protein was detected by immunofluorescence. Images were acquired by confocal microscopy. Representative transmitted light and sum projected images of NPC1-A488 and LDL-A546 are shown. Size bar = 25 μm.

**Figure S3.**
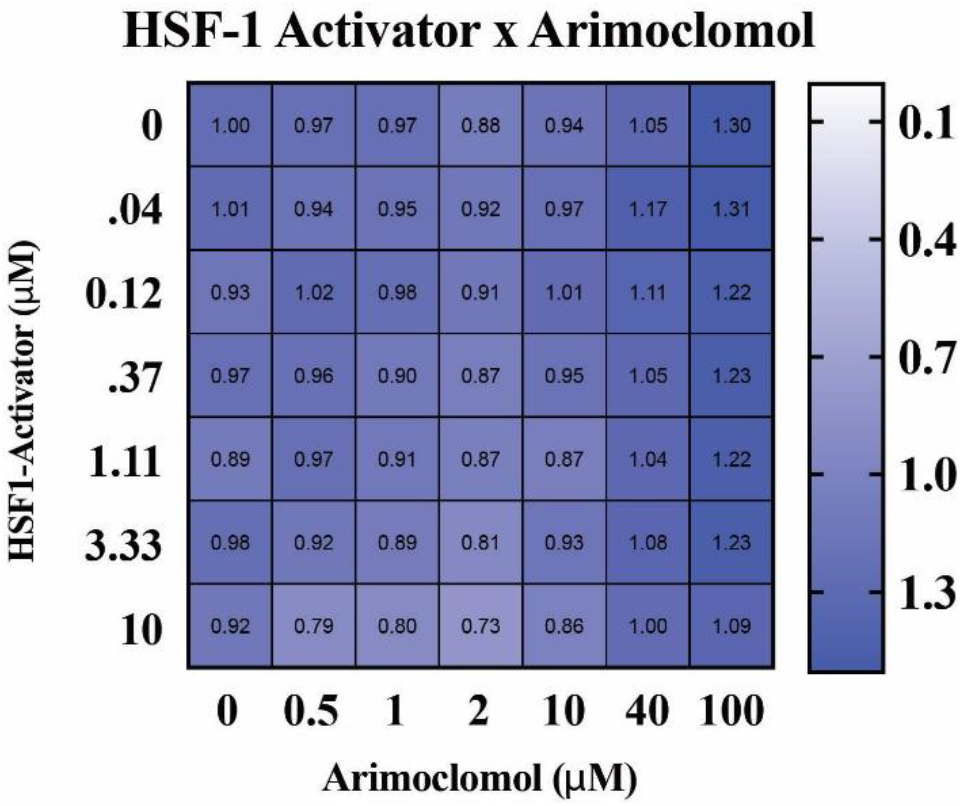
Cotreatment of HSF1a and arimoclomol. GM18453 fibroblasts were plated in 384-well plates and co-incubated with HSF1a and arimoclomol in a dose-dependent manner for 72 h. Heatmap value represents the average LSO of three independent experiments with four replicates per well per plate, each image had 100-300 cells, normalized to DMSO-treated cells. LSO values less than one represent a decrease in cholesterol storage, and values greater than one identify an increase in cholesterol storage. Concentration on vertical axis represents HSF1a and concentrations on horizontal axis is arimoclomol. All data are statistically insignificant.

